# Isometric exercise facilitates attention to salient events in women via the noradrenergic system

**DOI:** 10.1101/749002

**Authors:** Mara Mather, Ringo Huang, David Clewett, Shawn E. Nielsen, Ricardo Velasco, Kristie Tu, Sophia Han, Briana Kennedy

## Abstract

The locus coeruleus (LC) plays a critical role in regulating attention via the release of norepinephrine (NE), with levels of tonic LC activity constraining the intensity of phasic LC responses. However, the effects of manipulating tonic LC-NE activity on phasic activity have yet to be demonstrated in humans. In the current fMRI study, we used isometric handgrip to modulate tonic LC-NE activity in the time period immediately afterwards. During this post-handgrip time, an oddball detection task was used to probe how changes in tonic arousal influenced functional coordination between the LC and a right frontoparietal network that supports attentional selectivity. As expected, the frontoparietal network responded more to infrequent target and novel sounds than to frequent sounds. Across participants, greater LC-frontoparietal functional connectivity, pupil dilation, and faster oddball detection were all positively associated with LC MRI contrast from a neuromelanin-sensitive structural scan. Thus, LC structural integrity was related to LC functional dynamics and attentional performance during the oddball task. We also found that handgrip led to larger phasic pupil responses to oddball sounds, faster oddball detection speed, and greater frontoparietal network activation, suggesting that something that induces strong LC activity benefits attentional performance for at least the next few minutes. In addition, older women showed a similar benefit of handgrip on frontoparietal network activation as younger women, despite showing lower frontoparietal network activation overall. Together these findings suggest that a simple exercise may improve selective attention in healthy aging, at least for several minutes afterwards.

**Highlights:**

- We examined how handgrip affects arousal-attention dynamics during an oddball task
- Salient stimuli enhanced pupil dilation and were detected faster post-handgrip
- A decrease in tonic pupil size after handgrip suggests norepinephrine depletion
- MRI-assessed LC integrity was related to oddball detection and pupil dilation
- LC integrity also associated with LC-frontoparietal network functional connectivity

## 1. Introduction

The locus coeruleus (LC) is a small brainstem nucleus that integrates signals about arousal from a variety of brain systems and, via its widespread projections, releases norepinephrine in cortical regions to modulate processing. Under arousal, salient stimuli stand out even more than they would otherwise, while less salient stimuli are even more likely to be ignored, regardless of whether the salient stimuli are themselves emotionally arousing or not (e.g., Lee, Sakaki, Cheng, Velasco, & Mather, 2014; Sutherland & Mather, 2012). The LC seems to play a key role in this effect of arousal, in part by modulating frontoparietal attention network processes (Lee et al., 2018; Kennedy & Mather, in press; Mather, Clewett, Sakaki, & Harley, 2016).

In the current study, we examined LC-frontoparietal attention network dynamics during an oddball task. To test the influence of LC tonic activity on these dynamics, we randomly assigned half of the participants to complete a handgrip task just before the oddball task. Moreover, by including a structural MRI scan to assess LC integrity, eye tracking during the oddball task, and groups of participants of different ages with varying sex hormone levels, we were able to relate performance and LC-frontoparietal dynamics during the oddball task to LC structural integrity, pupil responses across the task, and individual differences related to LC function.

### 1.1. LC modulation of frontoparietal regions involved in salience detection

Frontal and parietal brain regions are among those most innervated by NE neurons from the LC (Brown, Crane, & Goldman, 1979; Gaspar, Berger, Febvret, Vigny, & Henry, 1989; Gatter & Powell, 1977; Kobayashi, Palkovits, Kopin, & Jacobowitz, 1974) and in postmortem human brain samples, norepinephrine levels are highest in frontoparietal regions (Javoy-Agid et al., 1989). Consistent with this, blood-oxygen-level dependent functional magnetic resonance imaging (BOLD fMRI) signal indicates that, during rest, slow oscillations in locus coeruleus activity are especially strongly correlated with oscillations in frontoparietal regions (Jacobs, Müller-Ehrenberg, Priovoulos, & Roebroeck, 2018; Zhang, Hu, Chao, & Li, 2016). Another indication of coordination between the LC and frontoparietal regions comes from a study in which pupil dilation reflecting task effort was associated with BOLD fMRI activity in regions overlapping with both the LC and the frontoparietal network (Alnæs et al., 2014).

Furthermore, the degree of influence the LC has on frontoparietal regions varies depending on levels of arousal or noradrenergic activity. For instance, administration of an α2 agonist modified frontoparietal functional connectivity differently during an attention task than during rest (Coull, Büchel, Friston, & Frith, 1999). Also, among younger adults, hearing a tone that predicted a possible electric shock increased activity in frontoparietal regions and increased functional connectivity between the LC and frontoparietal network regions compared with hearing a tone that predicted no shock (Lee et al., 2018). Of interest for the current study which includes post-menopausal women, older adults did not show as much increase in frontoparietal regions or in LC-frontoparietal functional connectivity under arousal as did younger adults (Lee et al., 2018).

Detecting salient oddballs is associated with LC activity (Aston-Jones, Rajkowski, Kubiak, & Alexinsky, 1994; Krebs, Park, Bombeke, & Boehler, 2018; Murphy, O’Connell, O’Sullivan, Robertson, & Balsters, 2014) as well as with both the ventral and dorsal aspects of frontoparietal attention networks involved in directing attention (Kim, 2014). Indeed, evidence suggests that LC-frontoparietal interactions increase the attentional impact of salient stimuli (Lee et al., 2018; Mather et al., 2016; Robertson, 2014). For instance, giving participants a β-adrenergic antagonist before they completed an oddball task reduced brain activity in response to oddballs in key frontoparietal attention network nodes (right ventrolateral prefrontal cortex and temporoparietal junction) (Strange & Dolan, 2007).

### 1.2. Influence of Tonic LC Activity on Phasic Responses

Elevated levels of tonic LC activity may constrain its potential for phasic activity (Gilzenrat, Nieuwenhuis, Jepma, & Cohen, 2010). For instance, recordings from monkey LC reveals that higher tonic LC activity is associated with episodes of greater distractibility and decreased LC phasic responses to target stimuli during attention tasks (Aston-Jones & Cohen, 2005). In the current study, we wanted to examine whether tonic levels of LC activity would influence phasic responses to oddballs. This is especially interesting in the context of aging. It is not known whether there are age differences in the levels of tonic LC activity. In general, older adults’ behavior is consistent with the Aston-Jones et al. profile of monkey behavior during a higher tonic LC state, as greater distractibility and inability to inhibit irrelevant information are hallmarks of aging (Hasher & Zacks, 1988; Kennedy & Mather, in press). For these reasons, we were interested in comparing brain activity during the oddball task at different tonic levels of arousal and LC activity.

Given that isometric handgrip elicits NE activity (e.g., Wallin et al., 1992; Wallin, Mörlin, & Hjemdahl, 1987) and modulates emotion-cognition interactions in the subsequent interval (Nielsen, Barber, Chai, Clewett, & Mather, 2015), it is likely to be modulating tonic LC activity during that interval. Engaging in an isometric handgrip task increases blood levels of NE (Lake, Ziegler, & Kopin, 1976; Vecht, Graham, & Sever, 1978) – and it does so more effectively than heavy dynamic work involving large muscles (Kozłowski, Brzezinska, Nazar, Kowalski, & Franczyk, 1973) or a stress task (Wallin et al., 1992). Handgrip also increases sympathetic motor nerve activity (Wallin et al., 1992) and pupil dilation (Nielsen & Mather, 2015; Nielsen et al., 2015), indicating involvement of the sympathetic system and likely LC engagement. Therefore, to examine how functional activity between the LC and frontoparietal brain regions is influenced by tonic levels of LC activity, we had half of the participants complete a handgrip task before the oddball detection task.

### 1.3. Assessing LC structure using magnetic resonance imaging (MRI)

Assessing the relationship between the LC and cognition in aging in living humans has been challenging due to the lack of landmarks to indicate the boundaries of the LC on typical structural MRI images. Fortunately for this endeavor, the LC has different magnetic properties than the surrounding brainstem and so yields hyperintense signal in specialized structural MRI sequences (Betts et al., 2019; Betts, Cardenas-Blanco, Kanowski, Jessen, & Düzel, 2017; Chen et al., 2014; Clewett et al., 2016; Dahl et al., in press; Keren et al., 2015; Keren, Lozar, Harris, Morgan, & Eckert, 2009; Hämmerer et al., 2018; Langley, Huddleston, Liu, & Hu, 2017; Liu et al., 2017; Priovoulos et al., 2018; Sasaki et al., 2006). It has not yet been fully determined which cellular properties of the LC lead to this hyperintense signal, but MRI structural contrast has been validated to correspond with the location of LC neurons in both humans and rodents (Keren et al., 2015; Watanabe, Tan, Wang, Martinez-Hernandez, & Frahm, 2019). In postmortem human brains, high contrast on T1 FSE scans corresponded with LC cells containing tyrosine hydroxylase (present in catecholamine neurons) and neuromelanin (a pigmented polymer that is sequestered in vesicles as a side effect of catecholamine metabolism) (Keren et al., 2015). In rodents, high contrast on magnetization transfer gradient-echo MRI scans was seen in the location of the LC in control animals but not in those genetically altered not to produce LC neurons (Watanabe et al., 2019). Thus, in late life, lower LC MRI contrast is likely to reflect fewer LC neurons due to neurodegeneration. This is consistent with findings that greater LC MRI contrast in older adults is associated with better memory and cognition (Clewett et al., 2016; Dahl et al., in press; Hämmerer et al., 2018). Furthermore, among younger adults, LC MRI contrast is associated with greater goal-relevant selectivity during memory encoding (Clewett, Huang, Velasco, Lee, & Mather, 2018), indicating that this structural indicator relates to LC function among younger adults as well as among older adults.

In the current study, we included a T1 FSE sequence to localize the LC and quantify its contrast. We then examined whether those with higher LC structural integrity based on this measure also showed stronger functional connectivity, or functional integration, between the LC and frontoparietal regions during a selective attention task known to engage both regions (e.g., Mccarthy, Luby, Gore, & Goldman-Rakic, 1997; Murphy et al., 2014).

### 1.4. Pupillometry as an indirect marker of LC activity

In addition to neuroimaging techniques, we acquired pupillometry data, as it reflects LC activity (Joshi, Li, Kalwani, & Gold, 2015; Reimer et al., 2016) and provides a sensitive metric of phasic responses to oddballs (Liao, Yoneya, Kidani, Kashino, & Furukawa, 2016; Murphy et al., 2014). Recordings of LC neurons in monkeys have found that LC neurons activate in response to infrequent target stimuli in visual oddball tasks, but respond only weakly or not at all to more frequent non-target items (for review see Nieuwenhuis, Aston-Jones, & Cohen, 2005). In humans, it has been challenging to assess LC activity directly on a trial-by-trial basis, but pupil dilation has been used as a proxy measure, as phasic LC stimulation leads to pupil dilation (Joshi et al., 2015; Reimer et al., 2016). Indeed, human studies indicate that oddball items produce greater pupil dilation than do frequent items (Gilzenrat et al., 2010; Kamp & Donchin, 2015; Murphy, Robertson, Balsters, & O’Connell, 2011). Furthermore, pupil activity during an oddball task is associated with blood-oxygen-level dependent (BOLD) activity in a dorsal pontine region overlapping with the location of the LC (Murphy et al., 2014). Because phasic LC responses are constrained by tonic levels of LC activity, we expected that the effects of handgrip would carry over to influence tonic and phasic pupil dynamics during the oddball detection task.

### 1.5. Potential Influences of Sex Hormones

Prior work suggests that sex hormone levels may influence LC function and how it interacts with frontoparietal attentional processes. In rats, the LC has both estradiol and progesterone receptors (Helena et al., 2006) and animal research also indicates that the LC is regulated by estrogen (Bangasser, Eck, & Sanchez, 2018; Herrera, Wang, & Mather, 2018). For instance, estradiol treatment of ovariectomized female monkeys and rats increases tyrosine hydroxylase and dopamine β-hydroxylase, both of which are critical for NE synthesis (Pau et al., 2000; Serova, Rivkin, Nakashima, & Sabban, 2002). In the current study, we compared older postmenopausal women not taking hormone supplements to two groups of younger women: one group in the luteal phase of their menstrual cycle and the other group on hormonal contraception. This design enabled us to examine whether any age differences might be related to group differences in hormone levels.

### 1.6. Overview

By having younger and older women engage in a handgrip task before an auditory oddball task during fMRI and eye-tracking, we were able to test three main hypotheses: 1) Changes in tonic LC activity induced by handgrip affects phasic responses to salient stimuli in the period immediately afterwards; 2) Higher structural LC MRI contrast is associated with greater LC functional connectivity with frontoparietal attention network regions; and 3) These markers of arousal and LC structure/function would relate to faster oddball detection. In addition, we examined whether the LC-frontoparietal dynamics changed with age and/or were associated with sex hormone status.

## 2. Methods

### 2.1. Participants

Ninety-two female participants were recruited from three groups: 1) younger women taking hormonal contraception, 2) younger women not taking hormonal contraception, in their luteal phase, and 3) older, postmenopausal women not taking hormone supplementation. Younger adult women were aged 18-33 years old (*M =* 21.2 years, *SD =* 2.89 years) and older adult women were aged 49-75 years old (*M =* 62.5 years, *SD =* 6.2 years). Participants were randomly assigned to one of two experimental conditions: handgrip and non-handgrip. We excluded some participants from all analyses due the following reasons: 1) One younger adult had an incomplete oddball functional scan; 2) One younger adult and two older adults had below chance performance on the oddball task; 3) One older adult did not make button responses during the oddball task. Among the remaining eighty-seven participants, there were fifteen participants in each of the six experimental groups except the luteal non-handgrip (n = 14), older adult non-handgrip (n = 14) and older adult handgrip (n = 14) groups. The study was approved by the University of Southern California’s Institutional Review Board and was carried out in accordance with the Code of Ethics of the World Medical Association. Younger adult participants received course credit or payment for their participation; all older women received monetary compensation.

In addition to the requirement that participants match one of the three hormone groups, inclusion criteria required that individuals be between 18-35 years old or 49-75 years old, a non-smoker, right-handed, understand English, have adequate hearing and vision to view computer and paper-pencil tasks (with correction if necessary), and be able to maximally squeeze a hand therapy ball of hard resistance with their right hand. Exclusion criteria included being left-handed, a history of cognitive impairment, current cigarette smoker, current use of beta-blocker or psychoactive medication, or failure to pass the magnetic resonance screening interview.

Participants were asked to refrain from caffeine, alcohol and cardiovascular exercise for the 12 hours prior to their experimental sessions to control for outside influences on baseline salivary alpha-amylase levels. To avoid contamination of salivary samples, participants were asked to fast one hour prior to each experimental session as well as refrain from brushing teeth and chewing gum within the hour before their appointment. Their compliance with these criteria was confirmed with them upon their arrival.

Younger adult participants were recruited based on their hormone status. All of the younger women were either naturally cycling with regular menstrual cycles or on monophasic hormonal contraception. The women who were naturally cycling were recruited in the “mid-luteal” phase of the menstrual cycle (18-24 days from the start of menstruation). We used a forward day count from the first self-reported day of menstruation to determine menstrual phase. Both sessions of the experiment were completed during the mid-luteal phase for all naturally cycling women.

For the women on monophasic hormonal contraception, all were on a combined formulation that had both ethinyl estradiol and a synthetic progestin. Women on birth control completed both sessions of the experiment between active pill days 7-21. One woman on hormonal contraception was excluded from final analyses because she was on triphasic hormonal contraception and failed to return for the second session. All post-menopausal women had not taken hormone replacement therapy within the last two years.

### 2.2. Materials and Procedures

All experimental sessions were conducted between 8:30 and 20:00 h. Prior to scanning, each participant completed informed consent, the magnetic resonance screening form, and the T2 authorization consent form. Participants completed a 50-minute scan session in a 3T Siemens Prisma^fit^ MRI scanner at the University of Southern California Dana and David Dornsife Cognitive Neuroscience Imaging Center. We used a 32-channel array coil for all scans.

#### 2.2.1. Structural Scans

The first two scans were a scout and then a brief T2 scan required for radiologist check for incidental findings. Next, we collected a high-resolution T1-weighted anatomic image for each participant (repetition time = 2,300ms, echo time = 2.26ms, inversion time = 1,060ms, flip angle = 9°, 176 sagittal slices, field of view = 256mm, bandwidth = 200 HZ/Px, voxel resolution = 1mm^3^ isotropic, and scan duration = 4 minutes and 44 seconds). After this, one neuromelanin sensitive-weighted MRI scan was collected using a T1-weighted FSE imaging sequence (repetition time = 750ms, echo time = 12ms, flip angle = 120°, 1 average to increase signal-to-noise ratio (SNR), 11 axial slices, field of view = 220mm, bandwidth = 287 HZ/px, slice thickness = 2.5mm, scan duration – 1 minute and 53 seconds).

#### 2.2.2. Assessing pupil dilation range

During the structural scans, we also collected maximum and minimum pupil size using the ASL long-range remote eye-tracker with a 60-Hz sampling rate. These data were collected using methods adopted from Piquado et al. (2010). Participants were presented with a black screen for 10s followed by a white screen for 10s (Piquado, Isaacowitz, & Wingfield, 2010).

#### 2.2.3. Handgrip task/control

After structural scans were completed, we administered a handgrip task in the scanner, with methods adapted from the isometric handgrip task employed by Topolovec et al. (2004) and Nielsen and Mather (2015). Participants were presented with a grayscale screen, with either a yellow or a blue circle (normed for luminance) in the center of the screen (Figure 1A). During the yellow circle “rest” periods, all subjects were instructed to relax with their palm on top of the ball. During the blue circle “grip” periods, handgrip participants were asked to pick up the hand therapy ball and squeeze it as hard as they could for five 18-s cycles. Control participants were instructed to pick up the ball but not to squeeze it. Participants were reminded to keep their eyes open and fixated on the cue throughout the session. Both conditions started with a 10-s yellow circle display to determine baseline pupil diameter, and pupil diameter was collected throughout the isometric handgrip or control ball hold task. Participants had the opportunity to practice their assigned task prior to entering the scanner.

**Figure 1.**
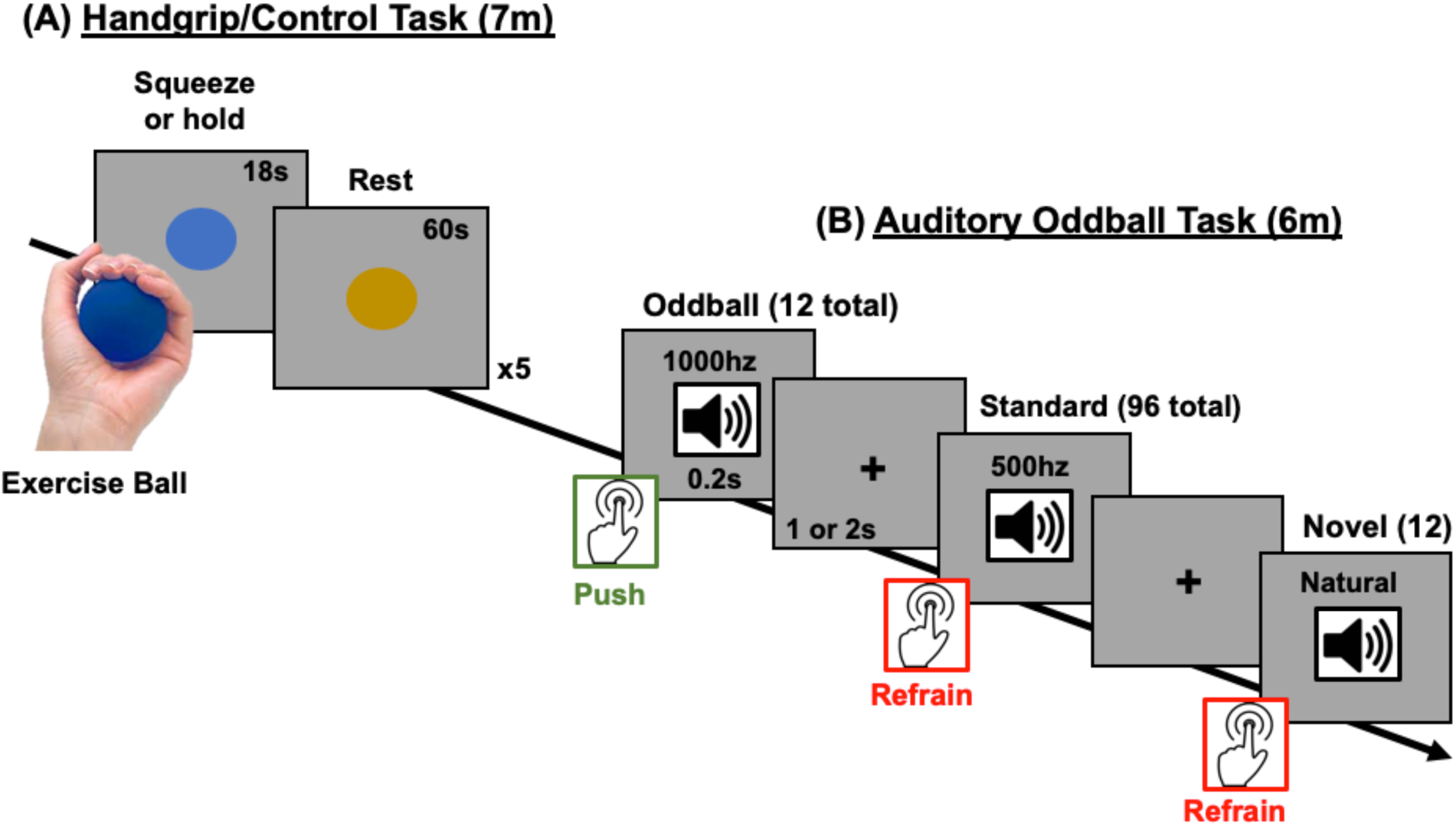
Experiment protocol for the (A) isometric handgrip task that was used to modulate tonic arousal and (B) the auditory oddball task that was used to assess salience detection. During the isometric handgrip task, participants squeezed (or, in the control group, just held) an exercise ball for 18s when a blue circle appeared and then rested for 60 seconds when a yellow circle appeared. There were 5 squeeze-rest cycles in total. Approximately 6-7 minutes after the beginning of the handgrip task, participants performed an oddball task in which infrequent high-pitched oddball tones (10% of trials), infrequent novel sounds (10% of trials), and frequent low-frequency tones (80% of trials) were presented in succession. Participants had to respond as quickly as possible via button press when they heard the oddball tone, but refrain from responding when they heard the novel and standard tones.

We used an echo-planar imaging sequence to measure BOLD fMRI signal during the task (repetition time = 2000ms, echo time = 25ms, flip angle = 90°, 1 average to increase signal-to-noise ratio, 41 axial slices, field of view = 192mm, bandwidth = 2520 HZ/px, slice thickness = 3mm, scan duration – 7 minutes and 4 seconds).

#### 2.2.4. Auditory oddball task

Immediately after the handgrip task, participants started the oddball detection task (Figure 1B). They heard a total of 120 tones throughout the approximately 6-min task. There were three different types of tones: 96 standard tones (500Hz), 12 novel tones (naturalistic sounds), and 12 oddball target tones (1000Hz). Each tone played for 200ms. Participants were instructed to press a button as quickly as possible when they heard the oddball tone, and to refrain from pressing the button when they heard a novel or standard tone. A jittered inter-stimulus interval of 1 or 2 seconds was inserted between each tone. During this time participants viewed a central fixation cross on a gray screen and could make their button response. The order of ISI durations was randomized. The same timings and stimulus order were used for every participant, which allowed use of a tensor independent component analysis to detect task-relevant functional brain networks (see Section 3.6).

Younger participants completed a second block of the oddball task but, due to time constraints, older participants did not complete this second block (older participants typically required longer than younger adults to get situated in the scanner and we were concerned about their tolerance of a long scanner session). To facilitate comparison across groups and maximize the impact of the preceding handgrip task, in the current analyses we only examined the first oddball block. We used an echo-planar imaging sequence to measure BOLD fMRI signal during the task (repetition time = 2000ms, echo time = 30ms, flip angle = 90°, 1 average to increase signal-to-noise ratio (SNR), 37 axial slices, field of view = 224mm, bandwidth = 2520 HZ/px, slice thickness = 3.5mm, scan duration – 6 minutes and 12 seconds).

#### 2.2.5. Timing of questionnaires and saliva samples

At the beginning of the experimental session, each participant rinsed their mouth by drinking an 8-oz. bottle of water. Next, they completed a demographic information packet, the PANAS, the CES-D, and the STAI. Approximately 10 min after their arrival, participants provided a 1-mL saliva sample using the “passive drool” collection method, and then entered the scanner.

After the oddball task, participants completed another task in the scanner for around 30 min (results not reported here) and then were removed from the scanner. At this point, they completed a second PANAS and a second 1-mL saliva sample via the “passive drool” method.

#### 2.2.6. Assays of saliva samples

Saliva samples were immediately frozen for a minimum of 24 hours to allow mucins to precipitate. Prior to the assays, they were thawed and centrifuged at 3,000 x g for 15 min to extract particulates from saliva. Clear supernatant was decanted into microtubes. The two saliva samples from the MRI session were then analyzed for salivary alpha-amylase (sAA), 17β-estradiol, and progesterone levels using Salimetrics, LLC (State College, PA) ELISA kits and measured optically using Molecular Devices, LLC SpectraMax M3 Multi-mode Microplate Reader (Sunnyvale, CA). From these samples, we determined the average levels of 17β-estradiol and progesterone hormones and used these averaged values for our hormone correlation analyses. Mean ± SEM values of 17β-estradiol and progesterone were within the expected ranges of the used assays for younger women (Salimetrics, LLC, State College, PA). For sAA, we considered just the baseline pre-scan value, as the sample at the end of the scan came over 30 min after completion of the oddball task and was likely influenced by arousal during the post-oddball task.

#### 2.2.7. Quantifying LC MRI contrast

We followed the methods employed in Clewett et al., (2016) to quantify LC MRI contrast as a proxy measure of LC structural integrity. Using the LC structural scan, two raters independently located the most inferior slice where the inferior colliculus was visible and then moved down two slices (7 mm), to a region where typically LC contrast is evident. Raters then placed a 3 x 3 voxel cross on the voxels with peak intensity. A dorsal pontine tegmentum reference 10 X 10 voxel ROI was then located 6 voxels more ventrally (i.e., further from the fourth ventricle, nearer the center of the brainstem) from whichever LC that was located more ventrally than the other, and equidistant between them in terms of right and left side of the body. If there were an odd number of voxels between the left and right LC ROIs, the reference ROI was placed one voxel closer to the left LC.

LC contrast-to-noise ratios were calculated by taking the difference between the LC and pontine tegmentum ROIs intensity values and dividing by the pontine tegmentum intensity. We then averaged the LC contrast-to-noise ratios obtained from each rater to obtain our LC contrast measure.

#### 2.2.8. Pupillometry preprocessing

Preprocessing of the pupillometry data was conducted using Matlab (Mathworks, MA). To remove blinks and other artifacts, we used a semi-automated program developed in our lab (github.com/EmotionCognitionLab/ET-remove-artifacts). First, each subject’s pupil timecourse was corrected with an automated blink-removal algorithm that detects blink events using the pupil timecourse’s velocity profile and then linearly interpolates over the detected regions. The details of the algorithm are described in depth elsewhere (Mathôt et al., 2013). Due to the high prevalence of non-blink noise in our recordings, several subjects had segments in their data that were undetected or improperly interpolated by the blink-removal algorithm. Therefore, a trained user either corrected the improperly interpolated segments or, if the duration of the noise exceeded one second, classified those segments as missing data.

To control for interindividual differences in pupil mechanics, we collected pupillary dark and light reflex data in response to black and white screens, respectively, during the structural scan prior to the handgrip and oddball tasks. However, there was a high number (N = 25) of participants with unusable noisy data while viewing the white screen. Thus, we calibrated the pupil signal using the data acquired during the dark screen period only. Each subject’s pupil data during the oddball task were normalized as a percentage of her maximum pupil diameter while viewing the dark screen using the following transformation formula, 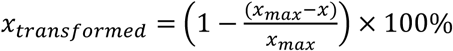, where *x_transformed_* is the new sample, *x* is the original sample, and *x_max_* is the max pupil diameter of the 500-ms window moving average when viewing the black screen. Effectively, this transformation accounts for individual differences in the pupil diameter ceiling as determined by the pupillary dark reflex mechanics.

For the pupillometry analyses of the handgrip task, we divided the pupil timecourse into five peri-cue-onset trials consisting of two trial events. The “Baseline” event consists of the 10 seconds prior to blue-cue onset, and the “Grip” event consists of the 18 seconds after the blue-cue onset. Trials were excluded from further pupil-related analyses if more than 50% of the preprocessed pupil signal in either the “Baseline” period or the “Grip” period were classified as missing data. If fewer than three trials survived the trial-level exclusion criteria, all of that participant’s data were excluded from any pupil-related analyses. After applying these exclusion criteria, 25 handgrip and 24 control subjects remained for the handgrip pupillometry analyses.

For the oddball pupillary analyses, we computed trial-by-trial baseline vs. change in pupil diameter to represent the tonic and phasic pupillary activity, respectively. Tonic activity is better captured by baseline pupil levels unaffected by the auditory stimuli, so our tonic pupil measure was the averaged pupil diameter of trial baselines, defined as the 1-s period prior to tone onset. We assessed phasic activity induced by the tone presentation in each trial by calculating the difference in mean pupil diameter between the 1-s post-tone offset period and the 1-s pre-tone onset period.

Trials were excluded from analyses if more than half of the samples in that trial were imputed with missing data indicators. Subjects were excluded from all pupil-related analyses if eye-tracking data or timing files was not collected for either the calibration or oddball task sessions. Subjects were also excluded from the tonic pupil analysis if more than 75% of total trials were missing and from the phasic pupil analysis if more than 75% of the trials in any of the three conditions were missing. Additionally, one subject was excluded from pupil analyses for the oddball task due to poor behavioral performance. After applying these exclusion criteria, 57 subjects (28 CTRL and 29 HG) remained for analysis of tonic pupil activity. One additional subject from this set did not meet criteria for analyses of phasic pupil activity, leaving 56 subjects (27 CTRL and 29 HG) for analysis of phasic pupil activity.

In a follow-up to the tonic pupil analysis, we segmented the pupil timecourse into three segments containing 40 trial each and performed the same analysis steps on each segment. The same subject-level exclusion criteria applied for the whole-timecourse analysis was used for each segment (i.e., subjects with more than 75% of total trials missing were excluded). Thus, for the follow-up tonic pupil analysis, 59 subjects (30 CTRL and 29 HG) remained for segment 1 analysis, 54 subjects (26 CTRL and 28 HG) remained for segment 2, and 51 subjects (24 CTRL and 27 HG) remained for segment 3.

## 3. Results

### 3.1. Handgrip task increased pupil size

A 3 (group: younger hormone, younger luteal, postmenopausal) X 2 (condition: handgrip, control) X 2 (event: baseline vs. ball hold) mixed-factor ANOVA on pupil diameter as a percentage of the individual’s maximal response to a dark screen revealed a main effect of event, *F*(1,43) = 65.11, *p* < .001, partial *η^2^* =.60 that was qualified by an interaction of event and condition, *F*(1,43) = 8.17, *p* = .007, partial *η^2^*=.16. While control participants showed larger relative pupil size during the ball holding phase (*M* = 71.66, *SE* = 1.50) than during the baseline rest phase (*M* = 69.11, *SE* = 1.62), handgrip participants showed a more extreme difference between the rest (*M* = 73.08, *SE* = 1.57) and ball hold phase (*M* = 67.73, *SE* = 1.70). There were no other significant effects (all *F* < 1).

Peri-cue-onset pupil timecourses for the handgrip and control groups are plotted in Figure 2. The dark green line below the two plots denotes regions where pupil diameter in the handgrip and control groups significantly differed (*p* < .01). Both group plots exhibited initial pupil dilation responses that reached similar peak magnitudes after the onset of the blue cue (t = 10s). At around t = 12.5s, both timecourses began to return to baseline levels, although the control group timecourse recovered more rapidly than the handgrip group. The handgrip and control plots did not significantly diverge until t = 15.2s, at which point the pupil dilation was more sustained in the handgrip compared to the control group. At the end of the “ball hold” period, marked by the onset of the yellow cue (t = 28s), another pupil dilation response is observed in both plots. However, unlike the response during “ball hold,” the post-dilation recovery for both groups occurred at similar rates.

**Figure 2.**
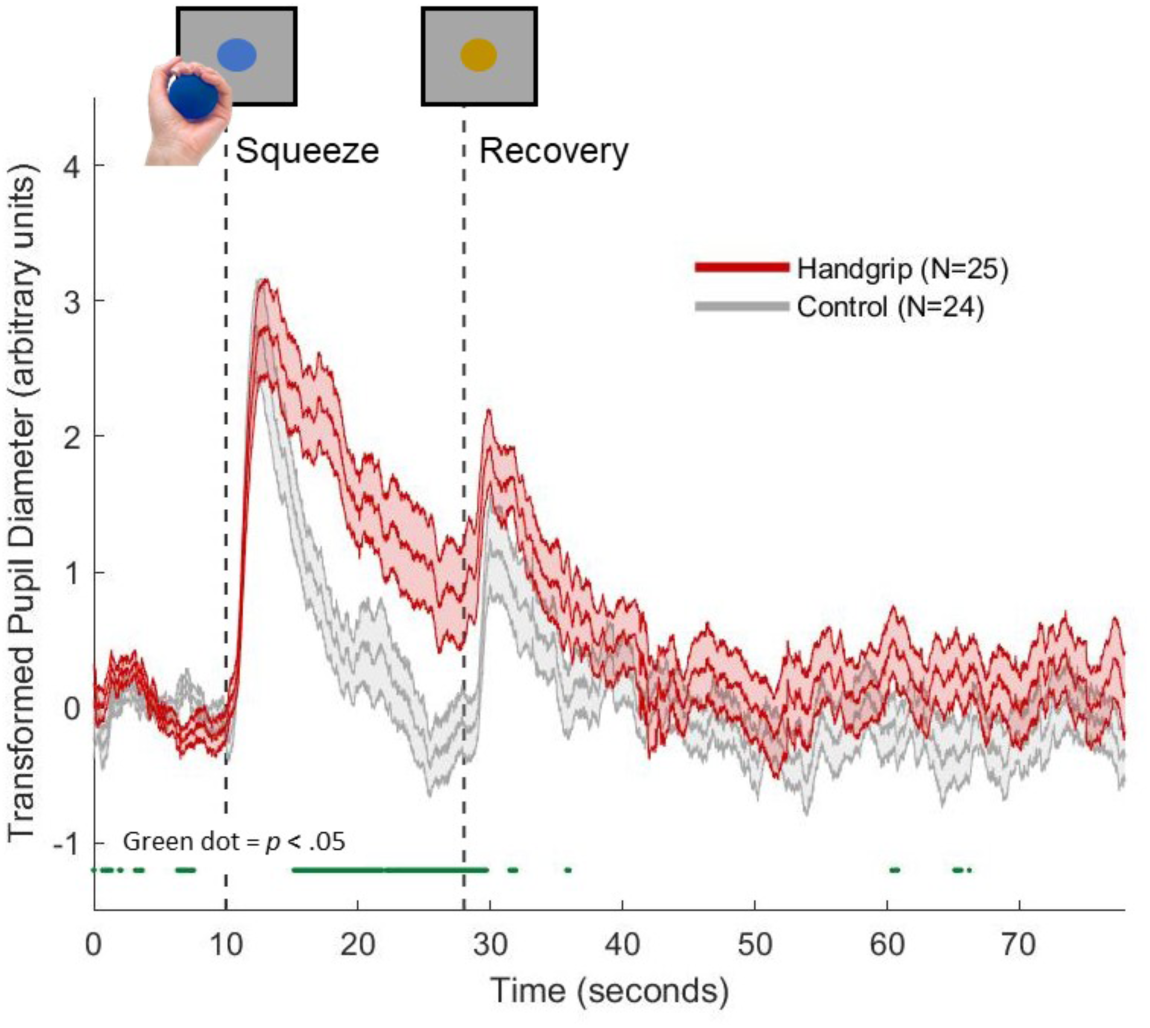
Trial-averaged pupil diameter during the isometric handgrip task for the handgrip group (red) and control group (grey). Green dots below the plots indicates samples with significant handgrip versus control group difference in pupil diameter (p < .05).

### 3.2. Behavioral performance during oddball task

As responses were only required for the oddballs, we examined corrected response rates for oddball trials. This was computed by subtracting the false alarm rate (mis-presses to standard or novel tones) from the oddball hit rate (correct button presses). A 2 (condition: handgrip, control) X 2 (group: younger hormone, younger luteal, postmenopausal) ANOVA revealed a main effect of group, *F*(2,81) = 4.87, *p* = .010, partial *η^2^* = .11. Post hoc t-tests indicated that this main effect was driven by postmenopausal women (*M* = .93, *SE* = .015) being significantly less accurate than women in the luteal group (*M* = .99, *SE* = .015; *p* = .026) and women in the hormonal contraception group (*M* = .99, *SE* = .014; *p* = .023). There was no significant interaction (*p* = .77).

Next, we examined response times for oddball tone trials, excluding trials where participants did not respond. A 2 (condition: handgrip, control) X 2 (group: younger hormone, younger luteal, postmenopausal) ANOVA revealed a main effect of group, *F*(2,81) = 3.22, *p* = .045, partial *η^2^* = .074. The younger luteal group had the quickest (*M* = 539 ms, *SE* = 16), the younger hormone group the next quickest (*M* = 568 ms, *SE* = 16), and postmenopausal women had the slowest responses (*M* = 599 ms, *SE* = 17). Participants in the handgrip condition were on average faster at detecting goal-relevant oddball tones (*M* = 550 ms, *SE* = 13) than those in the control condition (*M* = 588 ms, *SE* = 14), *F*(1,81) = 3.62, *p* = .048, partial *η^2^*= .047, indicating that handgrip has a carryover benefit on selective attention (Figure 3A). Post-hoc t-tests revealed that younger luteal women were significantly faster at responding to oddball tones than older postmenopausal women (*p* = .039). It is also noteworthy that the performance benefit of prior handgrip was marginally significant between the older women who had squeezed versus those who had not squeezed the exercise ball, *p* = .057. The interaction of condition and group did not achieve significance (*p* = .20).

**Figure 3.**
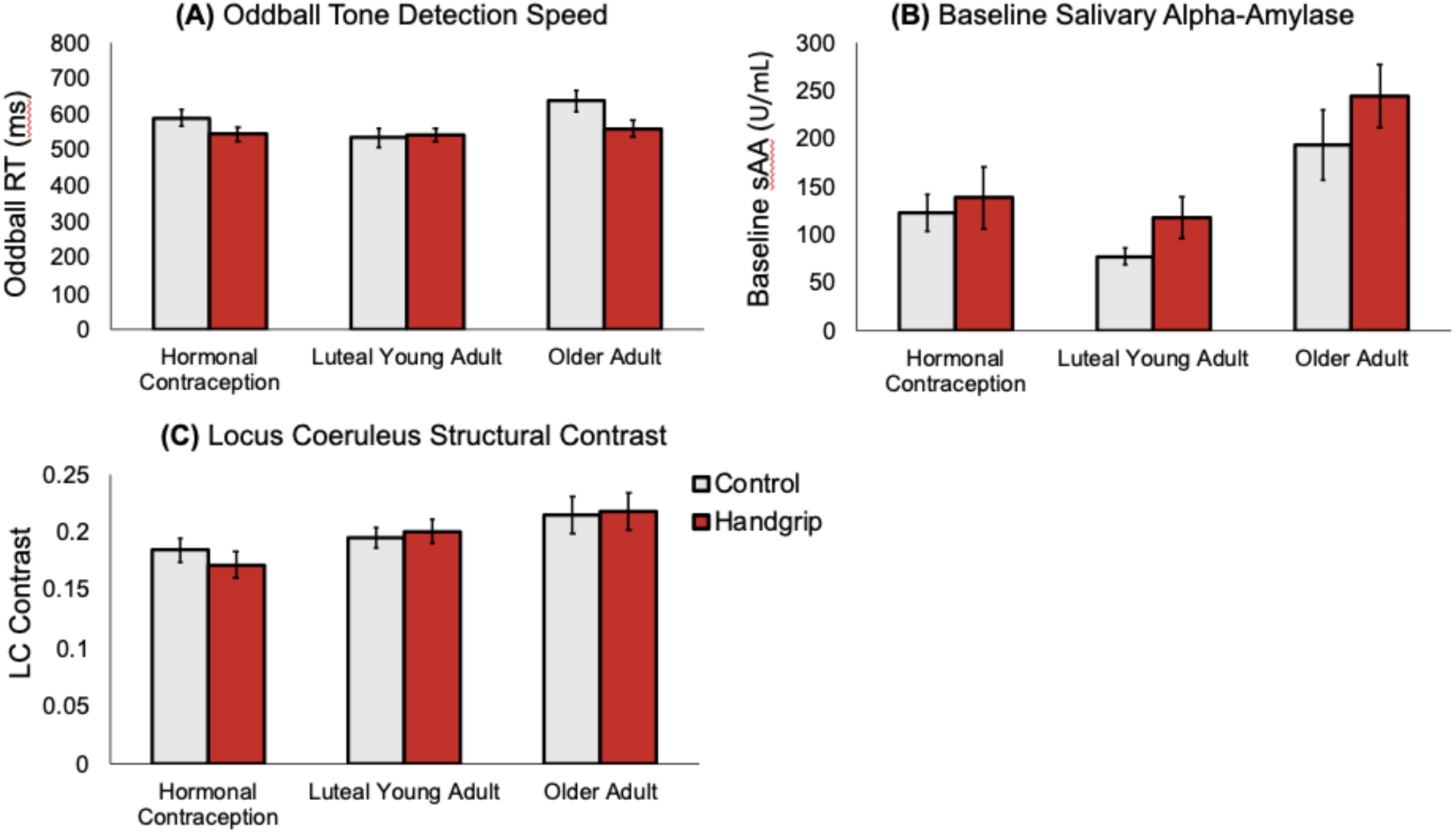
Between-group comparisons for (A) mean response times to oddball tones, (B) baseline levels of salivary alpha-amylase prior to MRI scanning, and (C) LC structural contrast.

### 3.3. Progesterone and estradiol levels

A 2 (condition: handgrip, control) X 3 (group: younger hormone, younger luteal, postmenopausal) ANOVA with estradiol level as the dependent variable revealed a significant effect of group, *F*(2, 80) = 12.86, *p* < .001, partial *η^2^* = .24. Post-hoc t-tests revealed that postmenopausal women had significantly lower estradiol levels than the younger luteal women (*p* < .001) and women on hormonal contraceptives (*p* = .007). There was a statistical trend towards younger luteal women having higher estradiol than women on hormonal contraceptives (*p* < .19). There were no interaction effects.

In a separate between-subjects ANOVA using progesterone level as the dependent variable revealed a significant main effect of group, *F*(2, 80) = 7.05, *p* = .002, partial *η^2^* = .15, and a marginally significant effect of condition, *F*(2, 80) = 3.92, *p* < .051, partial *η^2^* = .047. Post hoc t-tests revealed that younger luteal women had significantly higher progesterone levels than both younger women on contraceptives (*p* = .021) and the older postmenopausal women (*p* = .002). Thus, these analyses verified our hormone status groups by demonstrating that, at least for progesterone, luteal women had the highest levels of sex steroid hormones and both groups of younger women had higher sex steroid hormone levels than older postmenopausal women.

### 3.4. Salivary alpha amylase (sAA)

A 2 (condition: handgrip, control) X 3 (group: younger hormone, younger luteal, postmenopausal) ANOVA with baseline sAA as the dependent variable revealed a significant effect of group, *F*(2, 81) = 10.74, *p* < .001, partial *η^2^* = .21. As shown in Figure 3B, postmenopausal women had the highest baseline sAA values. Post-hoc t-tests revealed that the two younger groups did not differ significantly from each other (*p* = .63), but postmenopausal women had significantly higher sAA than the younger luteal women (*p* < .001) and women on hormonal contraceptives (*p* = .005). This increase in sAA with age is consistent with prior findings that older adults have a greater daily output of sAA than younger adults (Birditt, Tighe, Nevitt, & Zarit, 2017; Nater, Hoppmann, & Scott, 2013; Strahler, Berndt, Kirschbaum, & Rohleder, 2010).

### 3.5. Tonic and phasic pupillary activity during the oddball task

Figure 4 separately displays the group-averaged pupil diameter during the task sequence for those in the handgrip and control condition, with the timing of the oddball and novel tones indicated. To assess tonic activity, we performed a 2 (grip condition: handgrip, control) X 3 (hormone group: hormonal contraceptive, luteal phase, post-menopause) between-subjects ANOVA on baseline pupil diameter. No significant overall main effect of grip condition was observed, *F*(1,51) = 2.27, *p* = .14, partial *η*^2^ = .04. However, handgrip is likely to exert the greatest tonic difference early in the task. Thus, we segmented the session into 3 subsets containing 40 trials each and performed follow-up ANOVAs on baseline pupil diameter for each of the subsets. The analyses revealed a significant difference in baseline pupil size between the HG group (*M* = 67.77, *SE* = 1.73) and the CTRL group (*M* = 73.18, *SE* = 1.59), *F*(1,53) = 5.31, *p* = .03, partial *η*^2^ = .09, in the first subset, but not the other two. This suggests that the handgrip manipulation affected tonic activity in the first part of the subsequent task, possibly with weaker subthreshold effects in the later parts. No significant hormone group effects were found.

**Figure 4.**
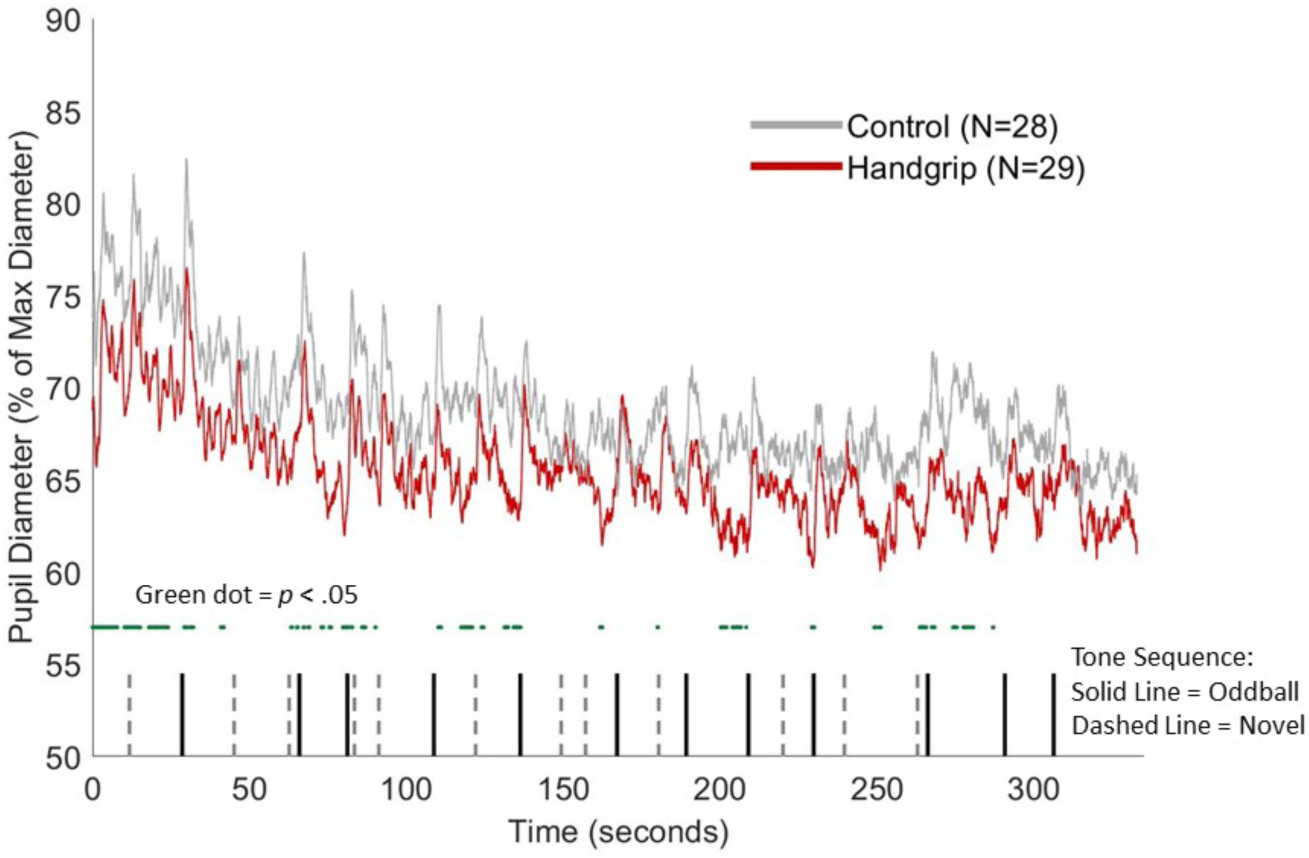
Group-averaged pupil diameter (as a percentage of the maximum diameter during the black-and-white screen baseline calibration task) during the oddball task sequence for the control group (grey) and handgrip group (red). The green dots below the plots indicate samples where the control pupil diameter is significantly greater than the handgrip pupil diameter (p < .05). Vertical black lines represent the onset timings of the goal-relevant oddball tones, whereas the dashed grey lines represented the onset of the novel sounds.

To assess phasic activity, we used the averaged change in pupil diameter in the oddball, novel and standard trials as the dependent variable in a 3 (tone type: oddball, novel and standard) X 2 (grip condition: handgrip, control) X 3 (hormone group: hormonal contraceptive, luteal phase, post-menopause) mixed-effects ANOVA. There was a strong effect of tone type, *F*(2,50) = 27.44, *p* < .001, partial *η*^2^ = .35, with the largest pupil dilation responses to oddball sounds (*M* = 1.51, *SE* = .20), next largest to novel sounds (*M* = .52, *SE* = .17) and smallest to standard sounds (*M* = .085, *SE* = .068). There was also a significant tone-type-by-grip condition interaction, *F*(2,50) = 3.31, *p* = .04, partial *η*^2^ = .06.

To follow up on this interaction, we ran separate 2 (grip condition: handgrip, control) X 3 (hormone group: hormonal contraceptive, luteal phase, post-menopause) between-subjects ANOVAs on pupil dilation response to oddball, novel, and standard tones. For the oddball tones, we found a significant main effect of grip condition on pupil dilation response, *F*(1,50) = 4.80, *p* =.03, partial *η*^2^ = .09, such that the dilation in the HG group (*M* = 1.95, *SE* = .28) was significantly greater than the dilation in the CTRL group (*M* = 1.07, *SE* = .29). In contrast, the ANOVAs did not show any significant differences on the pupil dilation response to novel and standard tones between the two grip conditions. Figure 5 displays the handgrip versus control group pupil dilation response to each tone type.

**Figure 5.**
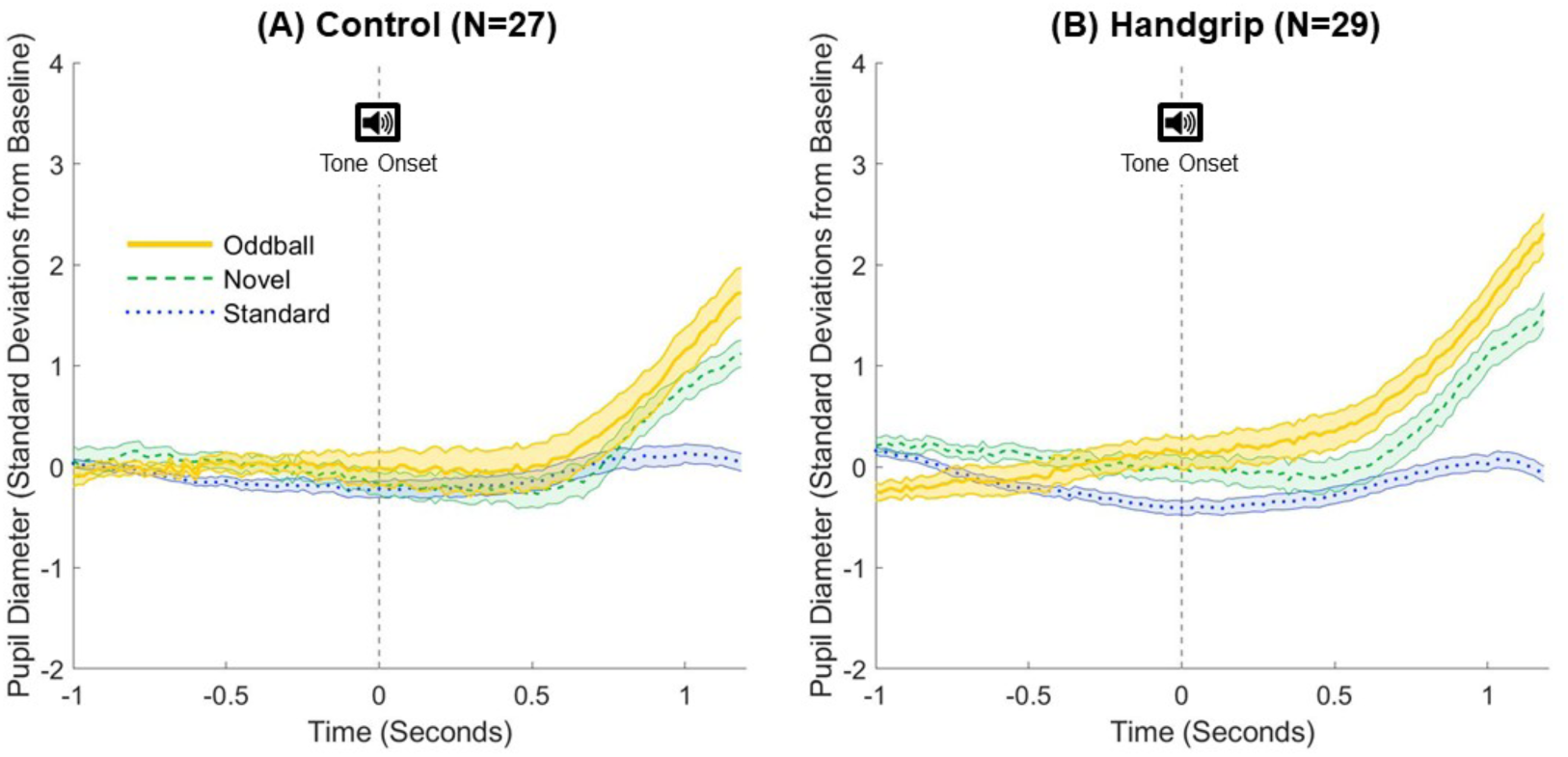
Trial-averaged phasic pupil responses to standard (blue dotted), novel (green dashed), and oddball (yellow solid) tones for the (A) control group and (B) handgrip group. Time t=0s (vertical dashed line) indicates the onset of the tone.

### 3.6. LC MRI contrast

A 2 (condition: handgrip, control) X 3 (group: younger hormone, younger luteal, postmenopausal) ANOVA with LC MRI contrast as the dependent variable revealed a significant effect of group, *F*(2,81) = 4.68, *p* = .012, partial *η^2^* = .10. As shown in Figure 3C, postmenopausal women had the highest LC MRI contrast values. Post-hoc t-tests revealed that the two younger groups did not differ significantly from each other (*p* = .33). Postmenopausal women had significantly greater LC contrast than the younger women on hormonal contraceptives (*p* = .009) but not the younger luteal women (*p* = .46). There were no significant interactions. The finding that older women had higher LC MRI contrast values than younger women is consistent with prior work using this fast spin echo sequence (Clewett et al., 2016).

### 3.7. Tensor independent component analysis to identify the frontoparietal network associated with oddball detection

All participants in our study received the same sequence of events in the oddball task. Thus, we were able to model a consistent timeline of events for all participants and employ this in a tensor independent component analysis (Beckmann & Smith, 2005) that provides a spatiotemporal decomposition and allows for contrasts between brain activity patterns during oddballs versus during other events. We constrained this tensor independent component analysis to 20 components in order to acquire well-established large-scale functional networks previously identified in the literature (Laird et al., 2011; Smith et al., 2009). We computed how much each of these 20 components were correlated to a template map of a previously identified right frontoparietal attention network (Fig 6A; independent component network #15 from Laird et al., 2011). We conducted the image correlation by using fslmaths (Jenkinson, Beckmann, Behrens, Woolrich, & Smith, 2012) to compute Z scores for each voxel within each image and then multiplying the two Z images and computing the mean value of the cross products (Mather, 2018).

**Figure 6.**
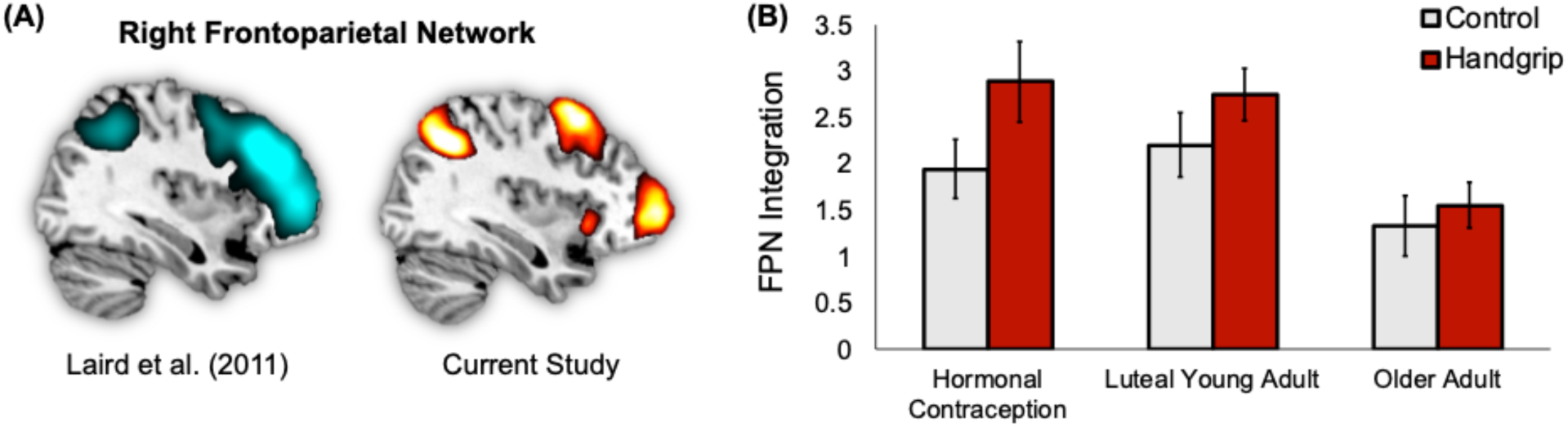
Right frontoparietal network connectivity during the oddball task. (A) A mask for the right frontoparietal attention network (FPN) from Laird et al., (2011) had a strong spatial correlation with component #4 identified by our tensor independent component analysis, a component that was most strongly associated with oddball events. (B) The handgrip condition increased participants’ loadings onto this right frontoparietal network component during the subsequent oddball task.

The component most strongly correlated to Laird et al.’s right frontoparietal network (*r* = .47) from our independent component analysis is shown in Figure 6A. To confirm that this component was best matched to the right frontoparietal network, we also examined the correlation between this component map and the other 19 networks Laird et al. (2011) had identified. No other correlations were as high (the next highest was *r* = .20).

This right frontoparietal network component had a strong full model fit, *F*(12,166) = 15.63, *p* < .001, meaning it was significantly related to oddball task events. Specific contrasts indicated this component was strongly related to both oddball, *z =* 7.03, *p <* .001, and novel, *z =* 4.86, *p* < .001, but not standard, *z =* 1.21, *p =* .11, sounds during the task. Both the oddball > novel contrast, *z =* 2.73, *p* = .0032, and the novel > standard contrast, *z =* 5.03, *p <* .001, were significant, as well as the oddball > standard contrast, z = 7.68, *p* < .001. Thus, this component responded most strongly to oddball targets, then to novel sounds, and least of all to standard sounds.

For the group level results comparing loadings on this component across participants, a 2 (condition: handgrip, control) X 3 (group: younger hormone, younger luteal, postmenopausal) ANOVA revealed main effects of handgrip condition and group (Figure 6B). Specifically, participants that completed the handgrip task (*M* = 2.43, *SE* = .19) loaded more onto this component during the task than did those who did not complete the handgrip task (*M* = 1.82, *SE* = .19, *F*(1,81) = 5.31, *p* = .024, *η^2^* = .062, indicating stronger functional connectivity in this network during the oddball task. The main effect of group, *F*(2,81) = 5.83, *p* = .004, *η^2^* = .13, emerged because postmenopausal women loaded less onto this component during the task sequence (*M* = 1.47, *SE* = .24) than did either the younger hormone contraception group (*M* = 2.42, *SE* = .23) or the younger luteal phase group (*M* = 2.48, *SE* = .23). There was not a significant interaction effect (*p* = .66).

### 3.8. LC-frontoparietal network connectivity analysis from dual regression

We took the individual spatial maps associated with the right frontoparietal attention network identified in the preceding analyses and used an LC mask to extract the Z-statistic estimates from this region-of-interest (ROI) for each participant to get an estimate of the functional connectivity of the LC with the right frontoparietal attention network during the oddball task. The LC mask was a probabilistic group mask derived based on the set of participants (N=74) with good neuromelanin scan registrations to MNI space (qualitatively determined by reviewing each FSE-to-MNI registration in FSLVIEW). The mask was thresholded at 20% probability, which means that at least 20% of the participants showed overlap between the voxels identified as having peak contrast for them and a given voxel in the group mask. A 2 (condition: handgrip, control) X 3 (group: younger hormone, younger luteal, postmenopausal) between-subjects ANOVA on the LC-frontoparietal connectivity scores for each participant yielded no significant main effects or interactions.

### 3.9. Associations between pupil dilation, LC contrast, LC-frontoparietal network functional connectivity, frontoparietal network loadings, and oddball reaction time

*Relationship with LC structural contrast.* Across groups, participants with greater structural LC MRI contrast had higher LC-frontoparietal network connectivity, *r*(85) = .25, *p* = .017 (Figure 7, top left panel). This correlation remained significant when age was partialled out, *r*(84) = .27, *p* = .011, as well as when both age and handgrip condition were partialled out, *r*(83) = .27, *p* = .012.

**Figure 7.**
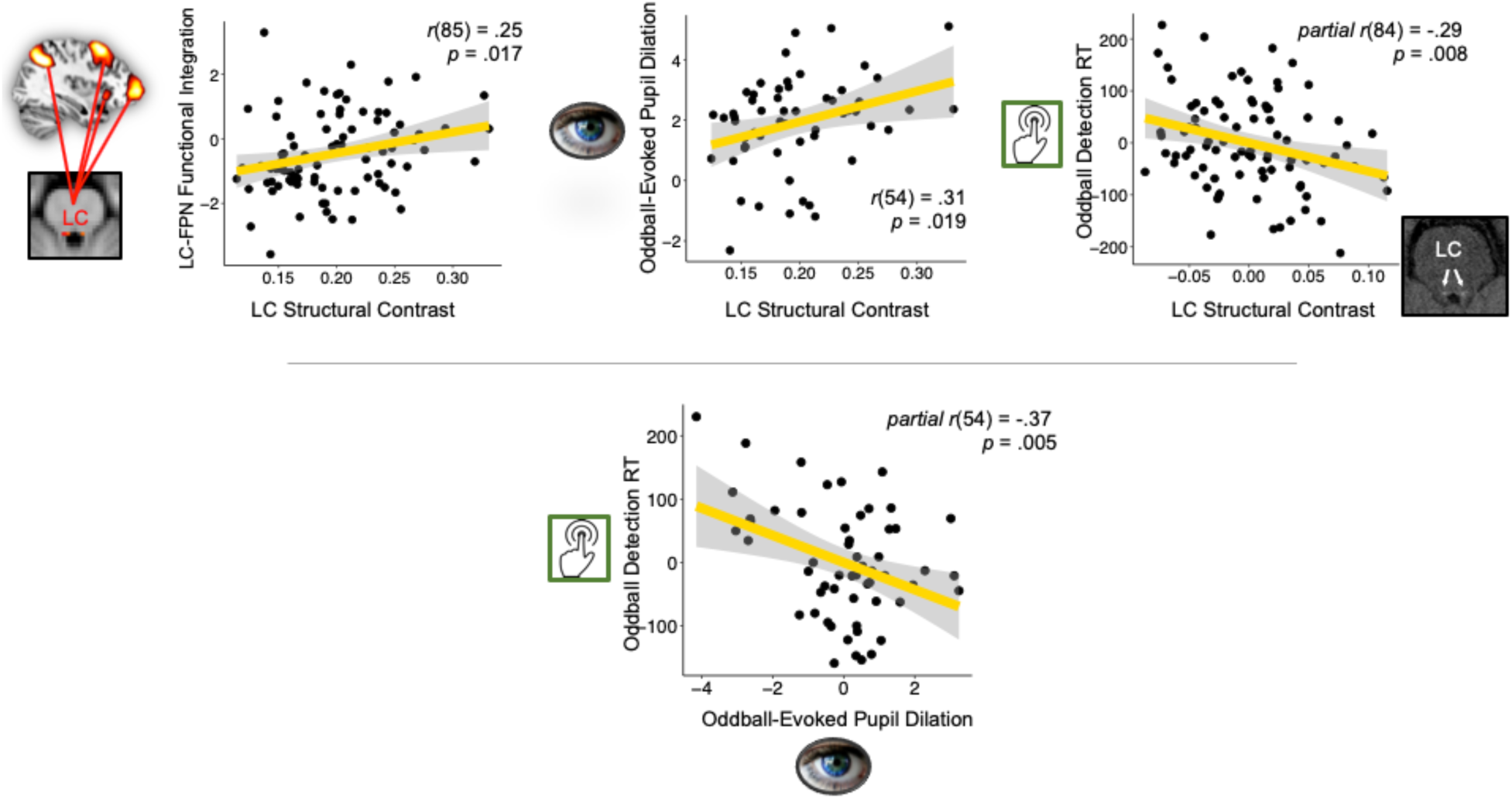
(Top panels) Associations between structural LC neuromelanin contrast-to-noise ratio and the LC-frontoparietal network (FPN) functional integration scores from the oddball task (left panel), oddball-evoked pupil dilations (middle panel), and residuals for oddball detection reaction times controlling for age (right panel). (Bottom panel) Association between the residuals for oddball-evoked pupil dilation and the residuals for reaction time to oddball tones controlling for the effects of age.

As accuracy on the task was near ceiling, we focused on response times to the oddball targets as a measure of performance. We partialled out age, as response time slowed with age. Across groups, participants with higher LC MRI contrast responded faster to the oddball targets, *r*(84) = -.29, *p* = .008 (Figure 7, top right panel). This correlation remained significant after controlling for handgrip condition as well, *r*(83) = -.30, *p* = .006.

*Relationship with pupil dilation.* We subtracted phasic pupil dilation to standard sounds from the phasic pupil dilation to oddball tones to dissociate oddball-evoked pupil dilations in response to goal relevance within each participant.

Across participants, greater oddball-evoked pupil dilations were associated with faster oddball detection (again controlling for age), *r*(54) = -.37, *p* = .005 (Figure 7, bottom panel). This relationship also remained significant after controlling for both age and condition (*p* < .05). Oddball-evoked pupil dilations were also positively correlated with LC MRI contrast, *r*(54) = 0.31, *p* = .019, which remained significant after controlling for age and after controlling for both age and condition (*p*’s < .05; Figure 7, top middle panel). Additionally, oddball-evoked pupil dilations were positively correlated with frontoparietal network loadings, *r*(54) = .28, *p* = .036. This relationship remained significant after controlling for age (*p* < .05), but became a trend after controlling for both age and condition (*p* = .13).

## Discussion

The locus coeruleus serves as the brain’s hub region for integrating signals about arousal (Mather, in press-b) and is believed to play a key role in modulating how focused attentional processes are on motivationally-relevant stimuli (Aston-Jones & Cohen, 2005; Mather et al., 2016; Nieuwenhuis, Gilzenrat, Holmes, & Cohen, 2005; Unsworth & Robison, 2017; Waterhouse & Navarra, 2019). Theoretical models propose that attentional selectivity for salient or high priority stimuli is particularly robust under conditions of increased phasic LC activity (e.g., Mather et al., 2016), which primarily emerge when tonic levels of LC activity are reduced (Aston-Jones & Cohen, 2005). However, there is only sparse empirical evidence of the brain and behavioral effects of modulating tonic arousal in humans. Here, we demonstrated that isometric handgrip reduced subsequent tonic levels of LC activity during an oddball task that involves noradrenergic system interactions with a right-lateralized frontoparietal attention network (Murphy et al., 2014; Strange & Dolan, 2007). Squeezing an exercise ball also led to larger phasic arousal responses to goal-relevant stimuli as well as better attentional performance. Additionally, we linked these beneficial effects of bursts in arousal to the structure and function of the LC-noradrenergic system, thereby underscoring the important role of this neuromodulatory system in supporting healthy attentional function across late adulthood.

Using a specialized fast spin echo MRI sequence, we found that LC MRI contrast, an in-vivo measure that has been linked to LC neuronal density (Keren et al., 2009), was associated with stronger functional integration between the LC and a right-lateralized frontoparietal attention network during an oddball detection task. Those participants with greater LC MRI contrast also responded faster to goal-relevant oddball stimuli and showed greater oddball-evoked pupil dilations, suggesting that the LC’s structural integrity constrains its ability to enhance selective attention. In a previous study, those with higher structural LC contrast showed greater selective LC activation during encoding of goal-relevant stimuli under threat (Clewett et al., 2018). Both those previous and the current findings indicate LC structure is related to its functional activity during selective attention tasks. Furthermore, our data suggest that the increased selectivity favoring highly salient or high priority information seen under arousal (Mather & Sutherland, 2011) is associated with LC structure across early and late adulthood.

In this study, we also examined whether sex hormones and age influenced dynamic interactions between the noradrenergic system and a frontoparietal attention network by testing three groups of women: younger women taking hormone contraception, naturally cycling younger women in their luteal phase, and post-menopausal women. As expected, compared with younger women, older women were slower and slightly less accurate at detecting salient oddball sounds. Consistent with this lower performance, postmenopausal women showed lower engagement of a right-lateralized frontoparietal network that responded to the oddball tones than did either of the two younger groups. Although animal research indicates that estradiol modulates LC activity (Bangasser et al., 2018; Herrera et al., 2018), our findings did not appear to be driven by hormone differences. The two groups of younger women with different hormone statuses did not show significant differences in frontoparietal network activation during the task. In addition, overall functional integration between the frontoparietal network and LC was similar across the three groups. However, a limitation of our study was that the N’s within subgroups of women (e.g., postmenopausal women who completed handgrip) were relatively small, limiting our ability to detect group differences. This was especially an issue for the pupil analyses, as not all participants had useable data.

Turning to the more robustly powered main effects of handgrip, there were some intriguing effects that suggest that squeezing an exercise ball affected subsequent tonic LC activity. First of all, functional integration within an oddball-sensitive frontoparietal network was greater during the oddball task among those who had just previously completed the handgrip than those who had not. Thus, the handgrip task affected frontoparietal network activity in the few minutes immediately afterwards. The pupil data suggest this effect was associated with reduced rather than greater tonic LC activity, as when compared to maximal pupil values assessed at the start of the session, pupil size was smaller in the post-handgrip condition during the first couple of minutes of the task. However, these pupil effects were not significant when examining the entire task time and so should be considered exploratory at this point. Phasic pupil responses to the oddball sounds were greater in the post-handgrip condition than in the no-handgrip condition, consistent with the notion that decreasing tonic LC activity can enhance phasic LC responses (Aston-Jones & Cohen, 2005; Gilzenrat et al., 2010).

It has long been known that isometric handgrip tasks lead to NE release (McAllister, 1979; Robertson et al., 1979), and recent findings confirm that pupils dilate during handgrip as well (Nielsen et al., 2015; Nielsen & Mather, 2015). However, little attention has been paid to how this strong burst of effort affects LC-NE activity during the period after the task. At first blush, our preliminary finding that handgrip leads to smaller pupil size, a putative marker of tonic LC activity, in the subsequent couple of minutes is somewhat surprising given that markers of arousal are elevated during handgrip itself. One intriguing possibility for these results, however, is that handgrip may have led to NE depletion for a few minutes after the task. Although currently a neglected area of research, in the 1960’s through the 1980’s, there were many animal studies documenting the phenomenon of NE depletion after arousing events. A variety of stressors including foot shock, exposure to responses of other animals receiving foot shock, exposure to an aggressor, cold stress, noise stimulation, isolation and immobilization lead to decreases in brain levels of NE afterwards in rats, mice, guinea pigs, and monkeys (Bliss, Ailion, & Zwanziger, 1968; Bliss & Zwanziger, 1966; Britton, Segal, Kuczenski, & Hauger, 1992; Glavin et al., 1983; Iimori et al., 1982; Maynert & Levi, 1964; Nakagawa, Tanaka, Kohno, Noda, & Nagasaki, 1981; Ordy, Samorajski, & Schroeder, 1966; Shinba, Ozawa, Yoshii, & Yamamoto, 2010; Tanaka et al., 1982; Weiss, Stone, & Harrell, 1970). This effect depends on an intact LC (Korf, Aghajanian, & Roth, 1973), and can be detected using in vivo microdialysis (e.g., Britton et al., 1992) as well as via post-mortem analyses. In rats, the degree of depletion shows a linear relationship with the duration of each shock as well as with the frequency of shocks, with depletion evident with as little as five minutes of intermittent shocks (Maynert & Levi, 1964). Confirming these older findings, sustained optogenetic activation of the LC also decreases subsequent NE levels throughout the brain (Zerbi et al., 2019). Our data suggest that, at least in humans, brief isometric exercise may also lead to NE depletion shortly thereafter. Replicating this exercise-induced NE depletion and determining its precise timing and its carryover effects on different aspects of cognition is an important direction for future research.

Our study adds to the growing literature on the relationship between the LC and attentional processing in the brain. We found that, across younger and older women, participants with higher structural LC MRI contrast also tended to show greater LC coordination with a frontoparietal network involved in detecting salient, goal-relevant information. Older women tended to show less engagement of a frontoparietal attention network during an oddball task. However, when older women performed an arousing handgrip task just beforehand, they showed increases in frontoparietal network integration that were comparable to younger adults in the no-handgrip condition. This finding suggests that a brief sustained episode of LC activity like that elicited during the five bouts of 18-s of maximal handgrip in our task may enhance subsequent attentional brain network function, leading to better selective attention. Interestingly, tonic pupil size decreased immediately after the handgrip, while phasic pupil responses to oddballs increased. This increase in phasic pupil responses was also related to faster detection of goal-relevant stimuli, suggesting that spikes in arousal facilitate adaptive responses to goal-relevant events.

These findings are consistent with the notion that older adults suffer from an overly high tonic level of LC-NE activity (Mather, in press-a; Weinshenker, 2018) and so may benefit from conditions that reduce tonic levels of NE. Future research is needed to test whether these findings replicate in males and whether relationships with sex hormones are seen with larger samples.

## References

Alnæs, D., Sneve, M. H., Espeseth, T., Endestad, T., van de Pavert, S. H. P., & Laeng, B. (2014). Pupil size signals mental effort deployed during multiple object tracking and predicts brain activity in the dorsal attention network and the locus coeruleus. Journal of Vision, 14(4).

Aston-Jones, G., & Cohen, J. D. (2005). An integrative theory of locus coeruleus-norepinephrine function: Adaptive gain and optimal performance. Annual Review of Neuroscience, 28, 403–450.

Aston-Jones, G., Rajkowski, J., Kubiak, P., & Alexinsky, T. (1994). Locus coeruleus neurons in monkey are selectively activated by attended cues in a vigilance task. The Journal of Neuroscience, 14(7), 4467–4480.

Bangasser, D. A., Eck, S. R., & Sanchez, E. O. (2018). Sex differences in stress reactivity in arousal and attention systems. Neuropsychopharmacology, 1.

Beckmann, C. F., & Smith, S. M. (2005). Tensorial extensions of independent component analysis for multisubject FMRI analysis. Neuroimage, 25(1), 294–311.

Betts, M. J., Cardenas-Blanco, A., Kanowski, M., Jessen, F., & Düzel, E. (2017). In vivo MRI assessment of the human locus coeruleus along its rostrocaudal extent in young and older adults. Neuroimage, 163, 150–159.

Betts, M. J., Kirilina, E., Otaduy, M. C., Ivanov, D., Acosta-Cabronero, J., Callaghan, M. F., … Passamonti, L. (2019). Locus coeruleus imaging as a biomarker for noradrenergic dysfunction in neurodegenerative diseases. Brain.

Birditt, K. S., Tighe, L. A., Nevitt, M. R., & Zarit, S. H. (2017). Daily social interactions and the biological stress response: Are there age differences in links between social interactions and alpha-amylase? The Gerontologist.

Bliss, E. L., Ailion, J., & Zwanziger, J. (1968). Metabolism of norepinephrine, serotonin and dopamine in rat brain with stress. Journal of Pharmacology and Experimental Therapeutics, 164(1), 122–134.

Bliss, E. L., & Zwanziger, J. (1966). Brain amines and emotional stress. Journal of Psychiatric Research, 4(3), 189–198.

Britton, K. T., Segal, D. S., Kuczenski, R., & Hauger, R. (1992). Dissociation between in vivo hippocampal norepinephrine response and behavioral/neuroendocrine responses to noise stress in rats. Brain Research, 574(1-2), 125–130.

Brown, R. M., Crane, A. M., & Goldman, P. S. (1979). Regional distribution of monoamines in the cerebral cortex and subcortical structures of the rhesus monkey: concentrations and in vivo synthesis rates. Brain Research, 168(1), 133–150.

Chen, X., Huddleston, D. E., Langley, J., Ahn, S., Barnum, C. J., Factor, S. A., … Hu, X. (2014). Simultaneous imaging of locus coeruleus and substantia nigra with a quantitative neuromelanin MRI approach. Magnetic Resonance Imaging, 32(10), 1301–1306.

Clewett, D., Lee, T. H., Greening, S. G., Ponzio, A., Margalit, E., & Mather, M. (2016). Neuromelanin marks the spot: Identifying a locus coeruleus biomarker of cognitive reserve in healthy aging. Neurobiology of Aging, 37, 117–126.

Clewett, D. V., Huang, R., Velasco, R., Lee, T.-H., & Mather, M. (2018). Locus coeruleus activity strengthens prioritized memories under arousal. Journal of Neuroscience, 38(6), 1558–1574.

Coull, J., Büchel, C., Friston, K., & Frith, C. (1999). Noradrenergically mediated plasticity in a human attentional neuronal network. Neuroimage, 10(6), 705–715.

Dahl, M. J., Mather, M., Duezel, S., Bodammer, N. C., Lindenberger, U., Kuehn, S., & Werkle-Bergner, M. (in press). Rostral locus coeruleus integrity is associated with better memory performance in older adults. Nature Human Behavior.

Gaspar, P., Berger, B., Febvret, A., Vigny, A., & Henry, J. P. (1989). Catecholamine innervation of the human cerebral cortex as revealed by comparative immunohistochemistry of tyrosine hydroxylase and dopamine-beta-hydroxylase. Journal of Comparative Neurology, 279(2), 249–271.

Gatter, K., & Powell, T. (1977). The projection of the locus coeruleus upon the neocortex in the macaque monkey. Neuroscience, 2(3), 441–445.

Gilzenrat, M. S., Nieuwenhuis, S., Jepma, M., & Cohen, J. D. (2010). Pupil diameter tracks changes in control state predicted by the adaptive gain theory of locus coeruleus function. Cognitive, Affective, & Behavioral Neuroscience, 10(2), 252–269.

Glavin, G. B., Tanaka, M., Tsuda, A., Kohno, Y., Hoaki, Y., & Nagasaki, N. (1983). Regional rat brain noradrenaline turnover in response to restraint stress. Pharmacology Biochemistry and Behavior, 19(2), 287–290.

Hämmerer, D., Callaghan, M. F., Hopkins, A., Kosciessa, J., Betts, M., Cardenas-Blanco, A., … Dolan, R. J. (2018). Locus coeruleus integrity in old age is selectively related to memories linked with salient negative events. Proceedings of the National Academy of Sciences, 115(9), 2228–2233.

Hasher, L., & Zacks, R. T. (1988). Working memory, comprehension, and aging: A review and a new view. The Psychology of Learning and Motivation: Advances in Research and Theory, 22, 193–225.

Helena, C. V. V., de Oliveira Poletini, M., Sanvitto, G. L., Hayashi, S., Franci, C. R., & Anselmo-Franci, J. A. (2006). Changes in α-estradiol receptor and progesterone receptor expression in the locus coeruleus and preoptic area throughout the rat estrous cycle. Journal of Endocrinology, 188(2), 155–165.

Herrera, A. Y., Wang, J., & Mather, M. (2018). The gist and details of sex differences in cognition and the brain: How parallels in sex differences across domains are shaped by the locus coeruleus and catecholamine systems. Progress in Neurobiology.

Iimori, K., Tanaka, M., Kohno, Y., Ida, Y., Nakagawa, R., Hoaki, Y., … Nagasaki, N. (1982). Psychological stress enhances noradrenaline turnover in specific brain regions in rats. Pharmacology Biochemistry and Behavior, 16(4), 637–640.

Jacobs, H. I., Müller-Ehrenberg, L., Priovoulos, N., & Roebroeck, A. (2018). Curvilinear locus coeruleus functional connectivity trajectories over the adult lifespan: a 7T MRI study. Neurobiology of Aging, 69, 167–176.

Javoy-Agid, F., Scatton, B., Ruberg, M., L’heureux, R., Cervera, P., Raisman, R., … Agid, Y. (1989). Distribution of monoaminergic, cholinergic, and GABAergic markers in the human cerebral cortex. Neuroscience, 29(2), 251–259.

Jenkinson, M., Beckmann, C. F., Behrens, T. E., Woolrich, M. W., & Smith, S. M. (2012). Fsl. Neuroimage, 62(2), 782–790.

Joshi, S., Li, Y., Kalwani, Rishi M., & Gold, Joshua I. (2015). Relationships between pupil diameter and neuronal activity in the locus coeruleus, colliculi, and cingulate cortex. Neuron, 89, 1–14.

Kamp, S. M., & Donchin, E. (2015). ERP and pupil responses to deviance in an oddball paradigm. Psychophysiology, 52(4), 460–471.

Kennedy, B., & Mather, M. (in press). Neural mechanisms underlying age-related changes in attentional selectivity. In G. R. Samanez-Larkin (Ed.), The Aging Brain. Washington, DC: American Psychological Association.

Keren, N. I., Lozar, C. T., Harris, K. C., Morgan, P. S., & Eckert, M. A. (2009). In vivo mapping of the human locus coeruleus. Neuroimage, 47(4), 1261–1267.

Keren, N. I., Taheri, S., Vazey, E. M., Morgan, P. S., Granholm, A.-C. E., Aston-Jones, G. S., & Eckert, M. A. (2015). Histologic validation of locus coeruleus MRI contrast in post-mortem tissue. Neuroimage, 113, 235–245.

Kim, H. (2014). Involvement of the dorsal and ventral attention networks in oddball stimulus processing: A meta-analysis. Human Brain Mapping, 35(5), 2265–2284.

Kobayashi, R. M., Palkovits, M., Kopin, I. J., & Jacobowitz, D. M. (1974). Biochemical mapping of noradrenergic nerves arising from the rat locus coeruleus. Brain Research, 77(2), 269–279.

Korf, J., Aghajanian, G., & Roth, R. (1973). Increased turnover of norepinephrine in the rat cerebral cortex during stress: role of the locus coeruleus. Neuropharmacology, 12(10), 933–938.

Kozłowski, S., Brzezinska, Z., Nazar, K., Kowalski, W., & Franczyk, M. (1973). Plasma catecholamines during sustained isometric exercise. Clinical Science, 45(6), 723–731.

Krebs, R. M., Park, H. R., Bombeke, K., & Boehler, C. N. (2018). Modulation of locus coeruleus activity by novel oddball stimuli. Brain imaging and behavior, 12(2), 577–584.

Laird, A. R., Fox, P. M., Eickhoff, S. B., Turner, J. A., Ray, K. L., McKay, D. R., … Fox, P. T. (2011). Behavioral interpretations of intrinsic connectivity networks. Journal of Cognitive Neuroscience, 23(12), 4022–4037.

Lake, C. R., Ziegler, M. G., & Kopin, I. J. (1976). Use of plasma norepinephrine for evaluation of sympathetic neuronal function in man. Life Sciences, 18(11), 1315–1325.

Langley, J., Huddleston, D. E., Liu, C. J., & Hu, X. (2017). Reproducibility of locus coeruleus and substantia nigra imaging with neuromelanin sensitive MRI. Magnetic Resonance Materials in Physics, Biology and Medicine, 30(2), 121–125.

Lee, T.-H., Greening, S. G., Ueno, T., Clewett, D., Ponzio, A., Sakaki, M., & Mather, M. (2018). Arousal increases neural gain via the locus coeruleus–noradrenaline system in younger adults but not in older adults. Nature Human Behaviour, 2(5), 356.

Lee, T. H., Sakaki, M., Cheng, R., Velasco, R., & Mather, M. (2014). Emotional arousal amplifies the effects of biased competition in the brain. Social Cognitive and Affective Neuroscience, 9(12), 2067–2077.

Liao, H.-I., Yoneya, M., Kidani, S., Kashino, M., & Furukawa, S. (2016). Human pupillary dilation response to deviant auditory stimuli: Effects of stimulus properties and voluntary attention. Frontiers in Neuroscience, 10, 43.

Liu, K. Y., Marijatta, F., Hämmerer, D., Acosta-Cabronero, J., Düzel, E., & Howard, R. J. (2017). Magnetic resonance imaging of the human locus coeruleus: A systematic review. Neuroscience and Biobehavioral Reviews, 83, 325–355.

Mather, M. (2018). MRI image correlation script. Retrieved from https://github.com/EmotionCognitionLab/FSL_Correlation

Mather, M. (in press-a). How arousal-related neurotransmitter systems compensate for age-related decline In A. Thomas & A. Gutchess (Eds.), Handbook of Cognitive Aging: A Life Course Perspective: Cambridge University Press.

Mather, M. (in press-b). The locus coeruleus-norepinephrine system role in cognition and how it changes with aging. In D. Poeppel, G. Mangun, & M. Gazzaniga (Eds.), The Cognitive Neurosciences. Cambridge, MA: MIT Press.

Mather, M., Clewett, D., Sakaki, M., & Harley, C. W. (2016). Norepinephrine ignites local hotspots of neuronal excitation: How arousal amplifies selectivity in perception and memory. Behavioral and Brain Sciences, 39, e200.

Mather, M., & Sutherland, M. R. (2011). Arousal-biased competition in perception and memory. Perspectives on Psychological Science, 6, 114–133.

Maynert, E., & Levi, O. (1964). Stress-induced release of brain norepinephrine and its inhibition by drugs. Journal of Pharmacology and Experimental Therapeutics, 143(1), 90–95.

McAllister, J. R. (1979). Effect of adrenergic receptor blockade on the responses to isometric handgrip: studies in normal and hypertensive subjects. Journal of Cardiovascular Pharmacology, 1(2), 253–263.

Mccarthy, G., Luby, M., Gore, J., & Goldman-Rakic, P. (1997). Infrequent events transiently activate human prefrontal and parietal cortex as measured by functional MRI. Journal of Neurophysiology, 77(3), 1630–1634.

Murphy, P. R., O’Connell, R. G., O’Sullivan, M., Robertson, I. H., & Balsters, J. H. (2014). Pupil diameter covaries with BOLD activity in human locus coeruleus. Human Brain Mapping, 35(8), 4140–4154.

Murphy, P. R., Robertson, I. H., Balsters, J. H., & O’Connell, R. G. (2011). Pupillometry and P3 index the locus coeruleus–noradrenergic arousal function in humans. Psychophysiology, 48(11), 1532–1543.

Nakagawa, R., Tanaka, M., Kohno, Y., Noda, Y., & Nagasaki, N. (1981). Regional responses of rat brain noradrenergic neurones to acute intense stress. Pharmacology Biochemistry and Behavior, 14(5), 729–732.

Nater, U. M., Hoppmann, C. A., & Scott, S. B. (2013). Diurnal profiles of salivary cortisol and alpha-amylase change across the adult lifespan: Evidence from repeated daily life assessments. Psychoneuroendocrinology, 38(12), 3167–3171.

Nielsen, S. E., Barber, S. J., Chai, A., Clewett, D. V., & Mather, M. (2015). Sympathetic arousal increases a negative memory bias in young women with low sex hormone levels. Psychoneuroendocrinology, 62, 96–106.

Nielsen, S. E., & Mather, M. (2015). Comparison of two isometric handgrip protocols on sympathetic arousal in women. Physiology and Behavior, 142, 5–13.

Nieuwenhuis, S., Aston-Jones, G., & Cohen, J. D. (2005). Decision making, the P3, and the locus coeruleus--norepinephrine system. Psychological Bulletin, 131(4), 510.

Nieuwenhuis, S., Gilzenrat, M. S., Holmes, B. D., & Cohen, J. D. (2005). The role of the locus coeruleus in mediating the attentional blink: A neurocomputational theory. Journal of Experimental Psychology: General, 134(3), 291–307.

Ordy, J., Samorajski, T., & Schroeder, D. (1966). Concurrent changes in hypothalamic and cardiac catecholamine levels after anesthetics, tranquilizers and stress in a subhuman primate. Journal of Pharmacology and Experimental Therapeutics, 152(3), 445–457.

Pau, K. Y., Hess, D., Kohama, S., Bao, J., Pau, C., & Spies, H. (2000). Oestrogen upregulates noradrenaline release in the mediobasal hypothalamus and tyrosine hydroxylase gene expression in the brainstem of ovariectomized rhesus macaques. Journal of Neuroendocrinology, 12(9), 899–909.

Piquado, T., Isaacowitz, D., & Wingfield, A. (2010). Pupillometry as a measure of cognitive effort in younger and older adults. Psychophysiology, 47(3), 560–569.

Priovoulos, N., Jacobs, H. I., Ivanov, D., Uludağ, K., Verhey, F. R., & Poser, B. A. (2018). High-resolution in vivo imaging of human locus coeruleus by magnetization transfer MRI at 3T and 7T. Neuroimage, 168, 427–436.

Reimer, J., McGinley, M. J., Liu, Y., Rodenkirch, C., Wang, Q., McCormick, D. A., & Tolias, A. S. (2016). Pupil fluctuations track rapid changes in adrenergic and cholinergic activity in cortex. Nature communications, 7, 13289.

Robertson, D., Johnson, G. A., Robertson, R. M., Nies, A. S., Shand, D. G., & Oates, J. A. (1979). Comparative assessment of stimuli that release neuronal and adrenomedullary catecholamines in man. Circulation, 59(4), 637–643.

Robertson, I. H. (2014). A right hemisphere role in cognitive reserve. Neurobiology of Aging, 35(6), 1375–1385.

Sasaki, M., Shibata, E., Tohyama, K., Takahashi, J., Otsuka, K., Tsuchiya, K., … Sakai, A. (2006). Neuromelanin magnetic resonance imaging of locus ceruleus and substantia nigra in Parkinson’s disease. NeuroReport, 17(11), 1215–1218.

Serova, L., Rivkin, M., Nakashima, A., & Sabban, E. L. (2002). Estradiol stimulates gene expression of norepinephrine biosynthetic enzymes in rat locus coeruleus. Neuroendocrinology, 75(3), 193–200.

Shinba, T., Ozawa, N., Yoshii, M., & Yamamoto, K.-i. (2010). Delayed increase of brain noradrenaline after acute footshock stress in rats. Neurochemical Research, 35(3), 412–417.

Smith, S. M., Fox, P. T., Miller, K. L., Glahn, D. C., Fox, P. M., Mackay, C. E., … Beckmann, C. F. (2009). Close correspondence of the brain’s functional architecture during activation and rest. Proceedings of the National Academy of Sciences of the United States of America, 106, 13040–13045.

Strahler, J., Berndt, C., Kirschbaum, C., & Rohleder, N. (2010). Aging diurnal rhythms and chronic stress: distinct alteration of diurnal rhythmicity of salivary α-amylase and cortisol. Biological Psychology, 84(2), 248–256.

Strange, B. A., & Dolan, R. J. (2007). Beta-adrenergic modulation of oddball responses in humans. Behav Brain Funct, 3, 29.

Sutherland, M. R., & Mather, M. (2012). Negative arousal amplifies the effects of saliency in short-term memory. Emotion, 12, 1367–1372.

Tanaka, M., Kohno, Y., Nakagawa, R., Ida, Y., Takeda, S., & Nagasaki, N. (1982). Time-related differences in noradrenaline turnover in rat brain regions by stress. Pharmacology Biochemistry and Behavior, 16(2), 315–319.

Topolovec, J. C., Gati, J. S., Menon, R. S., Shoemaker, J. K., & Cechetto, D. F. (2004). Human cardiovascular and gustatory brainstem sites observed by functional magnetic resonance imaging. Journal of Comparative Neurology, 471(4), 446–461.

Unsworth, N., & Robison, M. K. (2017). A locus coeruleus-norepinephrine account of individual differences in working memory capacity and attention control. Psychonomic bulletin & review, 24(4), 1282–1311.

Vecht, R., Graham, G., & Sever, P. (1978). Plasma noradrenaline concentrations during isometric exercise. Heart, 40(11), 1216–1220.

Wallin, B., Esler, M., Dorward, P., Eisenhofer, G., Ferrier, C., Westerman, R., & Jennings, G. (1992). Simultaneous measurements of cardiac noradrenaline spillover and sympathetic outflow to skeletal muscle in humans. The Journal of Physiology, 453(1), 45–58.

Wallin, B., Mörlin, C., & Hjemdahl, P. (1987). Muscle sympathetic activity and venous plasma noradrenaline concentrations during static exercise in normotensive and hypertensive subjects. Acta Physiologica Scandinavica, 129(4), 489–497.

Watanabe, T., Tan, Z., Wang, X., Martinez-Hernandez, A., & Frahm, J. (2019). Magnetic resonance imaging of noradrenergic neurons. Brain Structure and Function, 1–17.

Waterhouse, B. D., & Navarra, R. L. (2019). The locus coeruleus-norepinephrine system and sensory signal processing: A historical review and current perspectives. Brain Research, 1709, 1–15.

Weinshenker, D. (2018). Long road to ruin: noradrenergic dysfunction in neurodegenerative disease. Trends in Neurosciences.

Weiss, J. M., Stone, E. A., & Harrell, N. (1970). Coping behavior and brain norepinephrine level in rats. Journal of Comparative and Physiological Psychology, 72(1), 153.

Zerbi, V., Floriou-Servou, A., Markicevic, M., Vermeiren, Y., Sturman, O., Privitera, M., … Bohacek, J. (2019). Rapid Reconfiguration of the Functional Connectome after Chemogenetic Locus Coeruleus Activation. Neuron.

Zhang, S., Hu, S., Chao, H. H., & Li, C.-S. R. (2016). Resting-state functional connectivity of the locus coeruleus in humans: in comparison with the ventral tegmental area/substantia nigra pars compacta and the effects of age. Cerebral Cortex, 26(8), 3413–3427.

